# Electrostatic effects in proteins are governed by pH-redistribution of the conformational ensemble

**DOI:** 10.1101/2020.02.02.931253

**Authors:** Christos M. Kougentakis, Ananya Majumdar, E. Bertrand García-Moreno

**Affiliations:** T.C. Jenkins Department of Biophysics, Johns Hopkins University, Baltimore, MD; Graduate Program in Molecular Biophysics, Johns Hopkins University, Baltimore, MD; Biomolecular NMR Center, Johns Hopkins University, Baltimore, MD

## Abstract

The imperative for charges to be hydrated is one of the most important organizing principles in biology, responsible for the general architecture of biological macromolecules and for energy storage in the form of electrochemical gradients. Paradoxically, many functional sites in proteins have buried ionizable groups^1^. These groups are tolerated because they are usually buried in the neutral state^2^. However, when they become charged they can drive structural transitions to open states in which the charge can be stabilized, mostly through interactions with water^3^. This coupling between the ionization of a buried group and conformational reorganization is precisely the mechanism used by proteins to perform energy transduction^4,5,6^. By applying this principle to a family of 25 variants of staphylococcal nuclease with internal Lys residues, it was possible to characterize in detail the range of localized partial unfolding events that even a highly stable protein that unfolds cooperatively can undergo in response to H^+^-binding. Conformational states that constitute vanishingly small populations of the equilibrium native state ensemble of this protein were identified by correlation of structural and thermodynamic data, providing a map of the conformational landscape of this protein with unprecedented detail. The data demonstrate that the apparent p*K*_a_ values of buried ionizable residues are not determined by the properties of their microenvironment but by the intrinsic propensity of the protein to populate open states in which internal charged residues can be hydrated. The role of buried residues in functional sites in proteins relies on their ability to tune the conformational ensemble for redistribution in response to small changes in pH. These results provide the physical framework necessary for understanding the role of pH-driven conformational changes in driving biological energy transduction^4^, the identification of pH-sensing proteins in nature^7^, and for the engineering of pH-sensitive dynamics and function in *de novo* designed proteins^8^.

---

Buried ionizable residues in proteins are critical for many biochemical processes, especially energy transduction^4^, catalysis^1^, and pH-sensing^7^. Owing to the incompatibility of charges with hydrophobic environments, the p*K*_a_ values of these residues can be highly anomalous, shifted in the direction that promotes the neutral state^2^. In proteins such as ATP synthase^4^, cytochrome C oxidase^5^, and bacteriorhodopsin^6^, the key molecular events of the energy transduction step involve reorganization of the protein in response to the transient presence of a charge in a hydrophobic environment. This reorganization exposes the buried group to polar environments where their p*K*_a_ values are more normal^5^. It is well established that H^+^-coupled conformational changes require that proteins sample multiple conformations with different proton binding affinities (i.e., ionizable residues must have different p*K*_a_ values in the different conformational states)^9^. What is not understood is the nature of the conformational reorganization that can be coupled to the change in ionization state of a buried group. The study of 25 variants of staphylococcal nuclease (SNase) with Lys residues at internal positions^2^ presented here addresses this question directly.

The p*K*_a_ values of the internal Lys residues in SNase (Fig. 1A)^2^ were measured previously by linkage analysis of the pH-dependence of thermodynamic stability. In nineteen cases, the internal Lys p*K*_a_ is depressed relative to its p*K*_a_ in water, some by more than 5 pH units. Crystal structures show that the overall protein fold is unaffected when the buried Lys residues are neutral (Fig. S1)^10^. No correlation between either the structural features of the microenvironment of the buried Lys residues or the magnitude of the p*K*_a_ shifts is apparent. In a small number of cases, NMR spectroscopy and X-ray crystallography have provided evidence of structural reorganization in response to ionization of buried Lys residues, presumably to expose the group to bulk solvent^11^. The low throughput of these methods has previously prevented a systematic study of the types of conformational changes coupled to H^+^- binding to the buried Lys residues.

**Figure 1.**
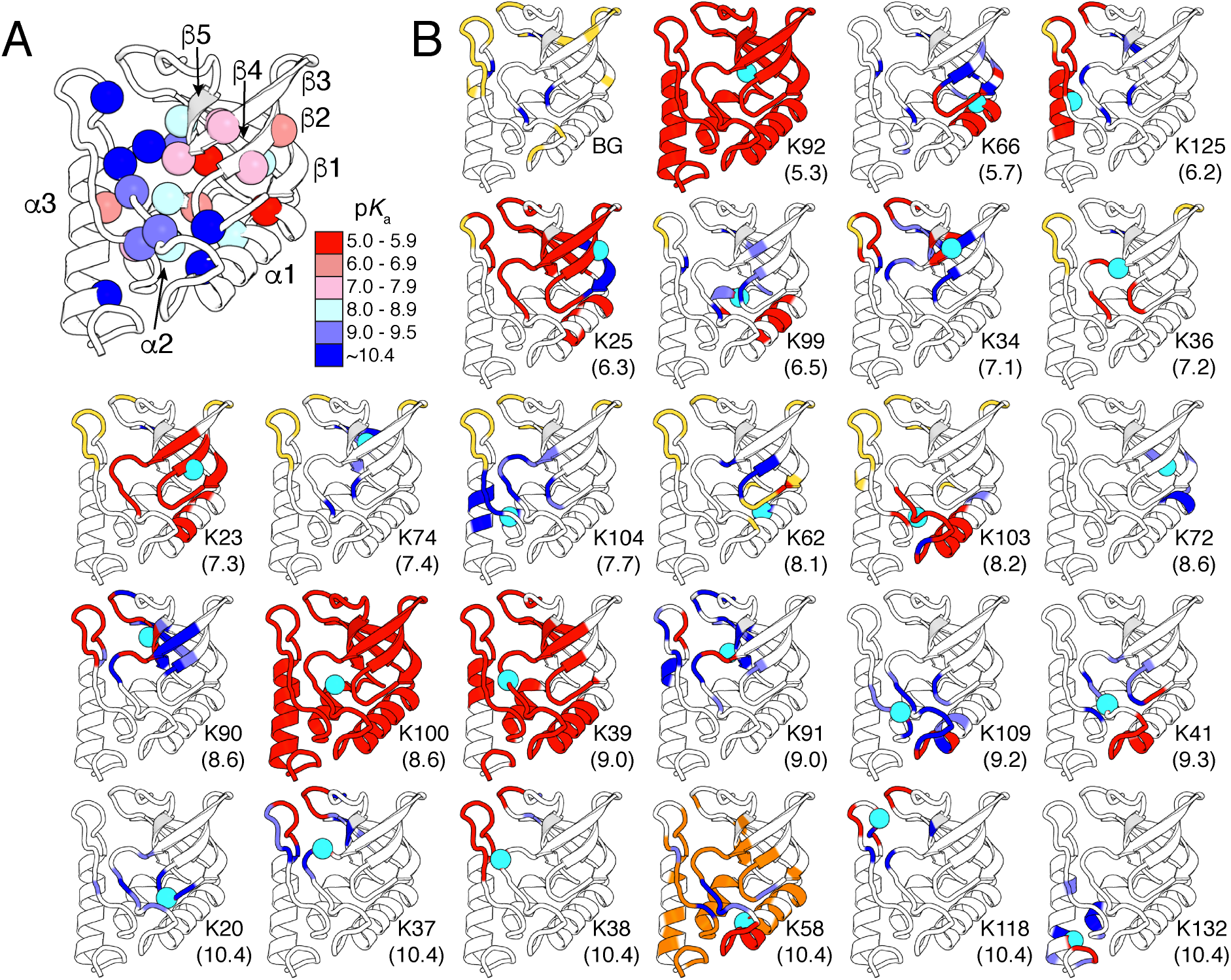
Structural reorganization coupled to the ionization of buried Lys residues. **(A)** Crystal structure of staphylococcal nuclease (PDB ID: 3BDC). Spheres identify sites of substitutions, color-coded according to the p*K*_a_ value of Lys residue at that position. **(B)** Chemical shift perturbations (CSPs) of backbone ^1^H_N_, ^1^Hα, ^13^Cα, ^13^Cα, ^13^C’, and ^15^N shifts. Background protein (BG) pH-dependent CSPs between pH 9.4 and 4.6 are shown for reference. For variants with internal Lys residues with p*K*_a_ values below 8.5, CSPs compare 1.2-1.5 pH units above and below the p*K*_a_ of the internal Lys. In order to avoid assignment difficulties at high pH due to base catalyzed exchange, CSP analysis for variants with p*K*_a_ values above 8.5 were calculated by comparing the variant with internal Lys at pH 7.2 with the background protein at pH 7.4. The p*K*_a_ values of the buried Lys are listed in parenthesis for each variant. Red denotes loss of resonance intensity, light and dark blue denote regions with moderate and large CSPs, respectively. Residues that were not observable owing to base catalyzed exchange at high pH are shown in yellow. Residues in slow exchange with two assignable conformations are shown in orange. A cyan sphere identifies the site of substitution with Lys.

Advances in data acquisition^12^ and analysis^13^ protocols have allowed for a dramatic increase in the throughput of NMR spectroscopy experiments, providing a means to describe with atomic detail the location and amplitude of structural reorganization that is coupled to the ionization of buried Lys residues in all 25 SNase variants. Chemical shift perturbations (CSP) of the backbone ^1^H, ^13^C, and ^15^N nuclei were measured for each variant at different pH values, or between the background and variants (Fig. 1). In two cases the ionization of the internal Lys triggered global unfolding (Fig. S2). In the other 23 variants the CSPs and/or loss of resonance intensities that were observed were consistent with local or subglobal changes in structure^14^ or dynamics^15^. In some instances, these changes were concomitant with an increase in random coil resonances (Fig. S3-26, Table S1-3). Structural analysis^16^ of the chemical shifts of variants at pH values where Lys is charged suggests that in many cases these changes reflect a loss of structure relative to the background protein or conformation at high pH (Fig. 2A). In other cases crystal structures confirm that the nature of the pH-dependent structural changes observed by NMR is partial unfolding (Fig. 2B-D)^11^. These data demonstrate unequivocally that ionization of the buried Lys residues is coupled to structural reorganization, shifting the population distribution from a predominantly fully folded (closed) state into an ensemble of locally, partially or fully unfolded (open) states in which the charged Lys residues are hydrated.

**Figure 2.**
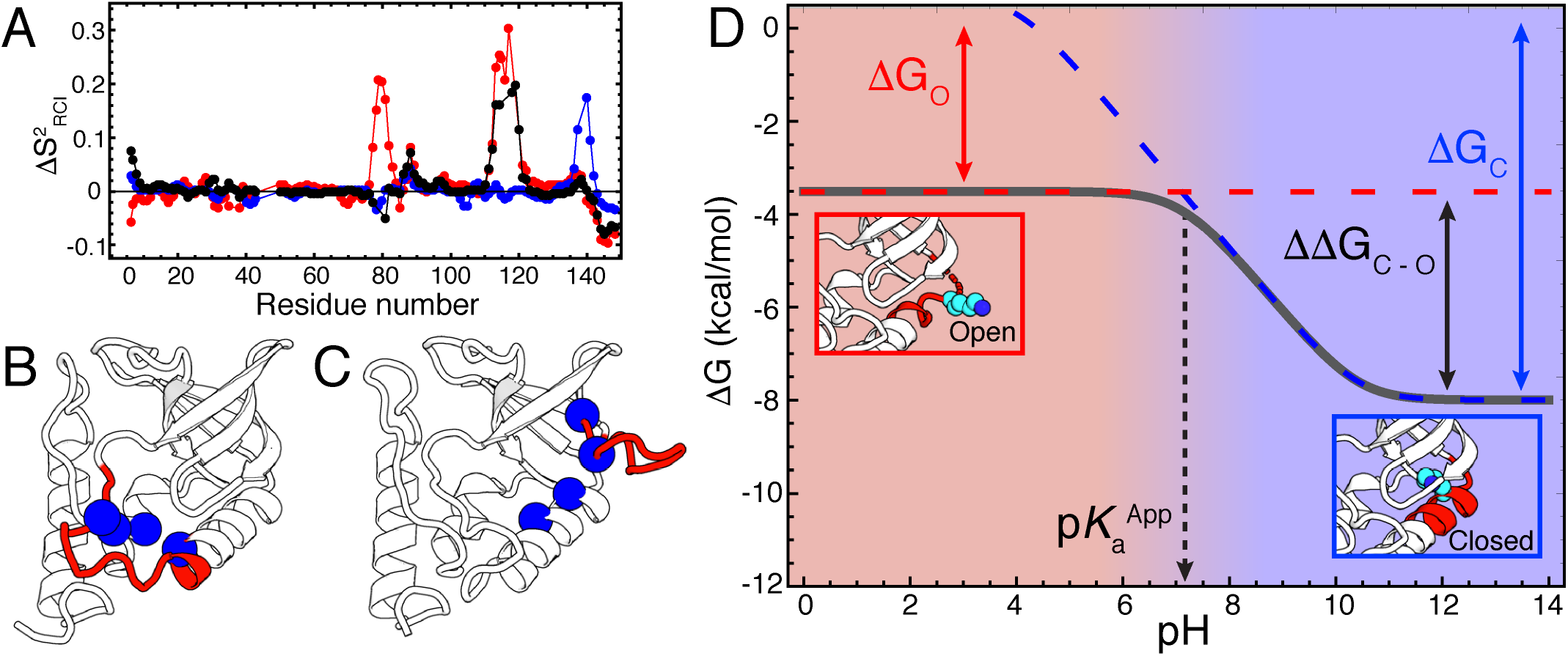
Structural and thermodynamic characterization of pH-dependent conformational changes necessary to hydrate buried Lys residues. **(A)** Difference in predicted order parameters (predicted using secondary shifts and the Random Coil Index) between the background protein at pH 7.4 and variants with K91 (red), K118 (black), and K132 (blue) at pH 7.2 (pH values where Lys is ionized). Positive values indicate regions that are predicted to be less structured in the variant relative to the background protein. Based on error estimates of the RCI analysis, values below 0.07-0.1 or from terminal residues should not be considered significant. **(B-C)** Crystal structures of variants with charged **(B)** R109 (PDB ID: 3D4W) and **(C)** E23 (PDB ID: 3TME). Regions showing evidence of structural reorganization relative to the background protein are highlighted in red. In Lys-containing variants where chemical shift changes or broadening are in regions showing partial unfolding as in the structures, the positions of the Lys substitutions are shown in blue spheres. **(D)** Simulation showing the pH-dependent thermodynamic stability of a protein with buried Lys. The solid gray line is simulated as the observed free energy difference between folded and unfolded states. ΔG_C_ is the free energy difference between the closed and unfolded states, which is pH-dependent below pH 10.4 (blue dashed line). ΔG_O_ is the free energy difference between the open and unfolded states (red dashed line). The energy gap between closed and open states is ΔΔG_C – O_, which is proportional to Δp*K*_a_. Closed and open states are represented by crystal structures of variants with K66 at pH 9 (closed, PDB ID: 3HZX) and pH 7 (open, PDB ID: 5CV5).

The thermodynamically determined p*K*_a_ values of these buried Lys residues are apparent p*K*_a_ values (p*K*_a_^App^) that do not report on the p*K*_a_ of the Lys in the buried state, but on the pH-dependent conformational equilibrium between the closed and the open states (Fig. 2D)^2,11^. In the open states the p*K*_a_ values should be comparable to the normal p*K*_a_ of Lys in water whereas in the closed states the p*K*_a_ must be lower than the measured p*K*_a_^App^. This observation has important implications for structure-based electrostatic calculations of p*K*_a_ values: the apparent p*K*_a_ values of buried groups are not determined by the electrostatic properties of the microenvironments around the buried ionizable moieties, but by the propensity of the protein to populate the open states.

To understand the physical origin of these pH-driven structural transitions, it is necessary to correlate the structural data from Figure 1 and 2 with equilibrium stability measurements. Owing to the shifted p*K*_a_ values of the buried Lys residues, at pH below the normal p*K*_a_ of 10.4 for Lys in water the stability of the closed states of each protein decrease in a pH-dependent manner (Fig. 2D). Although the open states are less stable than the closed states at high pH, their stability is pH-independent because in these states the Lys residues are exposed to water and their p*K*_a_ values are normal. As the pH decreases from values where Lys is neutral in water to where it is charged, the free energy gap between the closed and the open states decreases; p*K*_a_^App^ corresponds to the pH where this gap is zero.

At pH values below p*K*_a_^App^ the open state is the dominant state in solution (Fig. 2D), enabling structural characterization of the open state (with NMR spectroscopy) and measurement of its stability (with chemical denaturation experiments^2^). Because many variants showed similar patterns of CSPs or open state stabilities, it was possible to cluster variants based on similarity in the location or magnitude of their structural response. This allowed correlation between specific structural states populated by changes in pH with measured free energies (Fig. 3). The landscape shown two dimensionally in Fig. 3 shows that the region of SNase least destabilized by the ionization of buried Lys residues is the loop region between α-2 and α-3. (black in Fig. 3, black in Fig. 2A). Variants in which both this loop and the loop between α-4 and α-5 and in which the edge of α-3, α-4, and α-5 are perturbed (green in Fig. 3, red in Fig. 2A) are significantly less stable than the variants where only the loop between α-2 and α-3 appears to unfold. In both sets of variants ionization of Lys residues at three different positions triggered structural responses of similar magnitudes with similar open state stabilities. In contrast, the variants in which the turn between α-1 and α-2 or the loop between α-3 and α-1 are disrupted (orange and magenta in Fig. 3, Fig. 2B,C) exhibit greater variation in the character of the structural perturbations, and the associated stabilities of their open states.

**Figure 3.**
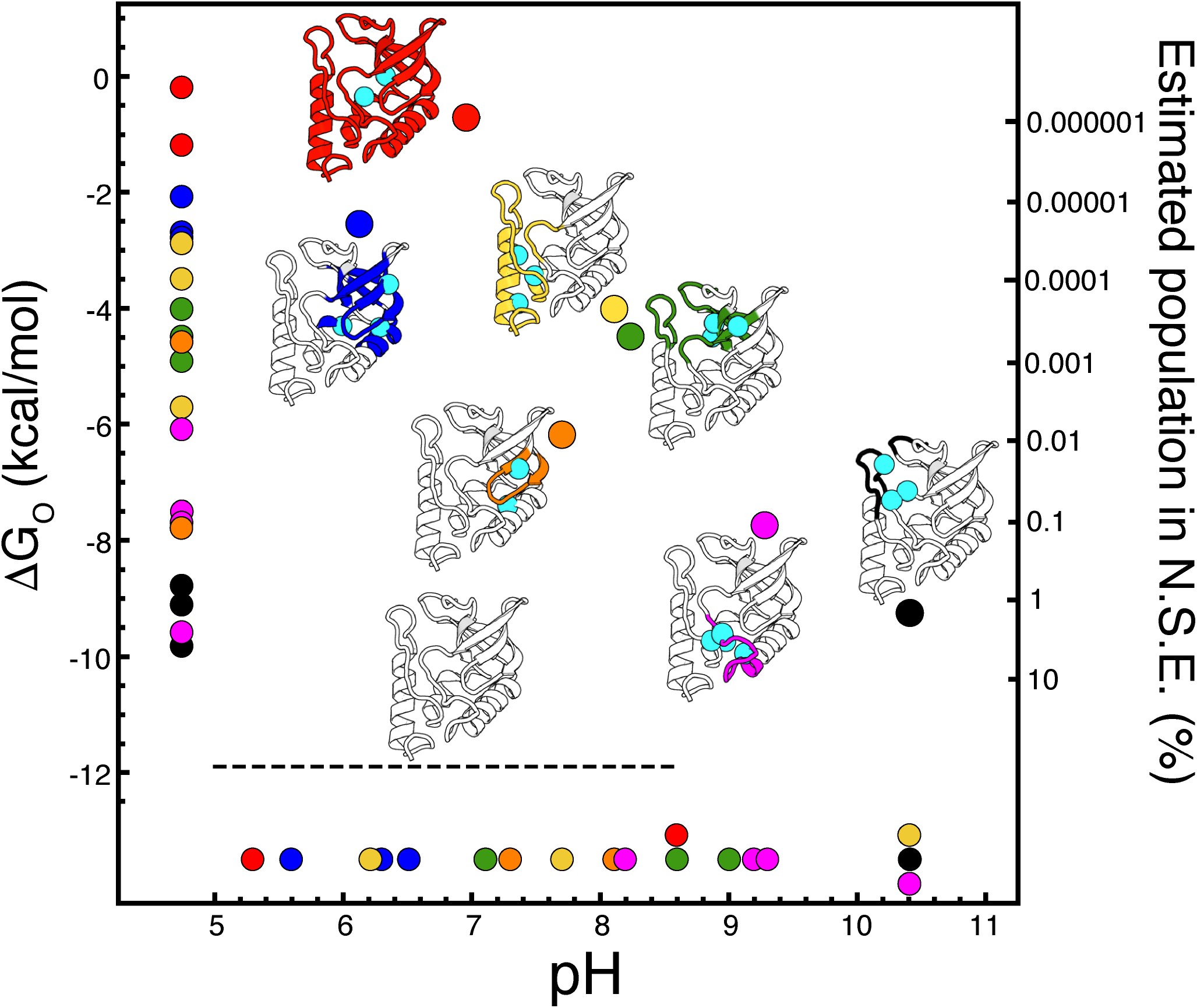
Conformational landscape of staphylococcal nuclease modulated by changes in pH. Correlation of the structural perturbations induced by the ionization of an internal Lys—detected with NMR spectroscopy—with thermodynamic stability measured by chemical denaturation for each protein under conditions of pH where the internal Lys is charged (small spheres, left axis). The p*K*_a_ values of the buried Lys residues are shown with small spheres on the bottom axis. The average open state stabilities and p*K*_a_ values are shown with large spheres. Structures in red represent variants with K92 and K100; blue with K25, K66 and K99; yellow with K104, K125 and K132; green with K34, K90 and K91; orange with K23 and K62; magenta with K41, K58, K103, and K109; and black with K37, K38, and K118. Cyan spheres identify the sites of substitution with Lys. The ground state in this diagram (bottom structure, black dotted line) corresponds to the thermodynamic stability of the background protein. All open state stabilities were made in a pH regime where stability of the background protein is pH-independent (11.8 kcal/mol, pH 5-8.5). The difference in stabilities between background protein and open states at identical solution conditions was used to estimate the approximate population of each state in the native state ensemble of this protein (right axis).

The least stable open states of SNase are those in which the two main subdomains of the protein, the OB-fold (blue in Fig. 3) and the interface made between α-2, α-3 and the α-barrel (yellow in Fig. 3), are significantly disrupted. Presumably the ionization of Lys residues buried in these regions disrupts important packing interactions. In the most destabilized variant in the yellow cluster (L125K) an increase in random coil resonances suggests partial unfolding of α-3. The variants in the blue cluster all show evidence of significant partial unfolding (^1^H-^15^N HSQC spectra in Fig. S4-7), with the α-barrel and α-2 affected to varying degrees; the C-terminus of α-1 is disrupted significantly in all of them. In all of these variants extensive disruption of the hydrophobic core of the OB-fold would be necessary to hydrate the buried Lys residue. Consistent with these observations, the blue regions have among the highest protection factors in hydrogen/deuterium exchange experiments^17^. Ionization of the buried Lys residue leads to global unfolding in only two variants (red in Fig. 3); in one case (N100K), fluorescence and circular dichroism spectroscopy provide evidence of some residual structure^2^. In five variants the structural responses could not be categorized either due to extensive resonance intensity loss in the native state or because the response is subtle and strictly local response.

The conformational substates that were identified in SNase by ionization of buried Lys residues are consistent with those identified previously with native state hydrogen deuterium exchange (Fig. S27)^17^, suggesting that the partially unfolded states in Figure 2 are relevant as low lying substates of the native state ensemble of this protein. The energy differences between the background protein and the open states allows for estimates of the population of these species in the native state ensemble of this protein. In most cases the substates could only exist at vanishingly small populations. These states are only populated to a significant extent by the truly local nature of these pH-driven structural changes. These partial unfolding reactions leave the rest of the cooperative folding core of the protein unaffected, as observed by the high m-values observed in chemical denaturation experiments^2^ (Fig. S28). Recent studies have shown that sparsely populated, partially unfolded states play important roles in tuning activity and allostery in well folded enzymes^18^. The results of this study are direct evidence that even well folded, stable proteins can access a wide range of partially unfolded states in response to relatively small changes in environmental conditions. The role of these conditionally disordered states in protein function has been underappreciated, given the difficulties in capturing these sparsely populated states with traditional structural biology methods. Although advances in structural methods have begun to access heterogeneity in the conformational ensemble^19^, this present study offers the most complete characterization of the native state ensemble of a protein.

Our data with SNase show that in the physiological range of pH^20^ a single ionizable residue with an anomalous p*K*_a_ can affect the stability and conformation of a protein in a significant way^2,21^. This suggests a mechanism whereby biological activity can be tuned through pH-dependent structural reorganization^22^, allosteric regulation^23^, and order-to-disorder transitions^24^. The role of buried residues is then to poise the conformational ensemble to respond to changes in pH, as well as other stimuli (e.g. photoillumination^25^, redox reactions^5^) that introduce charges in hydrophobic environments. With the large amount of structures available in the Protein Data Bank that can be mined with informatics approaches^7^, as well as the development of chemical probes that can selectively modify ionizable residues with anomalous p*K*_a_ values^26^, it should be possible to identify key residues and regions in proteins necessary for pH-sensing and regulation.

State-of-the-art electrostatics calculations, especially calculation of p*K*_a_ values, attempt to reproduce conformational reorganization coupled to the ionization of residues^27,28^. The data with SNase suggest two problems will have to be solved to increase the accuracy of these calculations. First, the calculations have to predict the structures of the open states, which will be challenging giving the highly local and disordered character of the reorganization (Fig. 1-2). Second, the Gibbs free energy differences between conformations will have to be calculated with high accuracy and precision, which will be difficult given the structural nuances described by Figure 3.

The conformation and stability of proteins has evolved to be sensitive to the chemical and physical properties of their environment. This is why proteins are endowed with the remarkable ability to respond to environmental changes, to interpret these changes as biological signals, and to amplify and transduce them into essential biological reactions. Understanding the mechanisms whereby pH regulates these properties of proteins is of special interest owing to increased recognition of pH as an essential biological signal^20^, and that pH dysregulation is observed in pathological states such as cancer^29^. This detailed structural and thermodynamic demonstration that a single ionizable residue can redistribute the conformational ensemble of a protein in response to small changes in pH in the physiological range has important implications for how proteins behave inside cells. Deeper understanding of the physical principles at play is necessary for improved mechanistic understanding not only of biological energy transduction^4^ and regulatory processes^20^ but also for improved prediction of pathological mutations that can make proteins sensitive to pH dysregulation^30^. Buried ionizable groups will also be useful to introduce pH-sensitive dynamics in engineered proteins by allowing them to transition between distinct conformational states, a hallmark of biological function^8^.

## Acknowledgements

Drs. Juliette Lecomte, Vincent Hilser, and Doug Barrick are thanked for discussions. Drs. Dominique Frueh, Scott Nichols and Bradley Harden assisted in the implementation of non-uniformly sampled pulse sequences. All data were acquired at the JHU Biomolecular NMR Center. This work was funded by NSF grant MCB1517378 and NIH grant GM061597 to B.G.M.E.

## Methods

### Protein purification and sample preparation

Bacterial stocks containing plasmids with single Lys substitutions introduced into a highly stable variant (deletion of a loop comprising residues 44-49, G50N, V51F, P117G, H124L, S128A, known as the Δ+PHS variant) of staphylococcal nuclease were previously prepared. Protein was prepared and stored as previously described^31^. For NMR experiments, proteins were thawed and buffer exchanged into 25 mM buffer and 100 mM KCl in 90% H_2_O/10% D_2_O. Reported pH values are the pH of buffer stocks protein was buffer exchanged into. Potassium acetate was used between pH 4-5.5, MES between pH 5.8-6.6, HEPES between pH 7-8, TAPS between pH 8-9, and CHES between pH 9-10. No buffer effects have been previously observed in SNase, and spectra collected in this study overlay with spectra from previous studies collected in different buffers or water^10^. Sample concentrations varied between 0.7-2.5 mM, usually from 1.0-1.5 mM. Variants with p*K*_a_ values between 5-8.5 were studied at two pH values corresponding to 1.2-1.5 pH units above and below the apparent p*K*_a_ value of the internal Lys residue (pH values for samples given in Fig. S3-14). All other samples were made at pH 7.2, and compared to a sample of the background protein at pH 7.4. Deviation of sample pH values after data collection from original buffer pH was observed to be less than 0.1 pH units.

### NMR Spectroscopy

All NMR experiments were performed on Bruker Avance or Avance II spectrometers equipped with cryogenic TCI probes. All data was collected at 25 °C. For backbone resonance assignments, 2D ^1^H-^15^N HSQC and 3D HNCACB, CBCACONH, HNCO, HBHACONH, CCC(CO)NH, and H(CCCO)NH spectra were collected. All spectra were collected with acquisition times of 6-8 ms in ^13^C (10-12 ms for HNCO), 8-10 ms in ^1^H and 23-24 ms in ^15^N in the indirect dimensions. Spectral widths in the indirect dimension for ^13^C were 62, 16, and 70 ppm for HNCACB/CBCACONH, HNCO, and CC(CO)NH spectra, respectively. Spectral widths in the indirect dimension for ^1^H were 8 and 11 ppm for HBHACONH and H(CCCO)NH experiments, respectively. The ^15^N spectral width was set to 25.5 ppm for all experiments. Carriers were adjusted based on spectra, but were generally 116-119 in ^15^N, 42 ppm for aliphatic ^13^C spectra and 177-178 ppm for the HNCO experiment. ^1^H carrier was consistently on the water frequency (4.76 ppm when accounting for salt concentration). 8 scans were collected per FID for each experiment except the CCCONH and H(CCCO)NH experiments, for which 16 scans per FID were required for adequate signal to noise, and the HNCO, for which only 4 scans per FID were necessary. All 3D experiments were collected using non-uniform sampling procedures with schedules generated using the Poisson Gap sampler developed by the Wagner lab^12^. Experiments were sampled at 15%, except for the CCCONH and (H)CCCONH, which were sampled at 17.5%. Spectra were reconstructed using NESTA-NMR^32^. Spectra were processed using NMRPipe^33^ and analyzed using NMRFAM-Sparky^34^.

### Assignment procedure

The ^1^H-^15^N HSQC, HNCACB, CBCACONH, and HNCO were manually peak picked. The automated assignment protocol of PINE-SPARKY was used to rapidly assign manually picked peaks^13,35^. In general, accuracy of this protocol was 90-100%, except in cases where broadened broke inter-residue connectivities, in which accuracy could drop to 50-60%. Spectra were then manually assigned to correct misassigned resonances from PINE, and to assign the ^1^Ha resonances using the HBHACONH and H(CCCO)NH spectra. Side chain patterns in the CC(CO)NH and H(CCCO)NH spectra were used to confirm backbone assignments from CBCACONH and HNCACB spectra, though no side chains past the Cα carbon were assigned for this study. All spectra underwent at least one more round of manual assignment to prevent misassignment of resonances.

### Chemical shift perturbation analysis

For the 12 variants for which data was collected at two pH values, the CSPs were calculated by subtracting the low pH data from the high pH data set. The CSPs were evaluated based on the CSPs observed between pH 9.4 and 4.6 in the background protein. This range of pH covers the entire range of pH values used in this study, and accounts for the titration of multiple residues (Fig. S3). The Cα resonances of Asp-21, His-121, and His-8 were excluded from the analysis of Cα CSPs as their large pH dependent CSPs reflect a change of charge state of an adjacent moiety, and significantly skew the standard deviation. Residues in the variants that showed at least two atoms having CSPs more than two standard deviations from the mean of the background protein CSPs were marked as having large CSPs (light blue), while residues with at least two atoms having CSPs more than three standard deviations from the mean of the background protein were marked as having very large CSPs (blue). This cutoff was chosen because the only residues that met the criteria for large or very large CSPs in the background protein were titrating residues (His-8, Asp-21, His-121) and one residue that is hydrogen bonded to the side chain of Asp-21 (Thr-41). Residues were identified as having intensity loss if the resonances for more than 3 out of 6 of the backbone nuclei are broadened beyond identification below the apparent p*K*_a_ value of the buried Lys. If this was only observed at pH values above the apparent p*K*_a_ value of the buried Lys residue, this was assumed to be due to base catalyzed exchange at high pH values necessary to deprotonate the buried Lys residue. Residues for which less than 3 chemical shifts could be measured for different reasons (e.g., multiple resonance overlap) were excluded from this cutoff. Mapping structural changes using weighted chemical shifts did not change the distribution of CSPs shown in Fig. 1.

### Structural studies of alternative states

Crystal structure of background protein and variants with neutral Lys were solved in previous studies^31,10^. Crystal structures of alternative states (A109R, V23E, and V66K) were solved in previous studies^36,37,11^. The V23E and V66K variants required secondary mutations to crystallize (D21N, T33V, T41I, S58A for V23E, K64G and E67G for V66K). Structural information was obtained through analysis of the 6 backbone chemical shifts measured in this study. TALOS-N was used to for torsion angle and secondary structure prediction^38^. Predicted order parameters (S^2^_RCI_) were generated using the RCI algorithm^16^, to assess changes in local dynamics.

### Simulation of thermodynamic stabilities

The thermodynamic stability of a protein with a buried Lys residue was simulated as^2^:

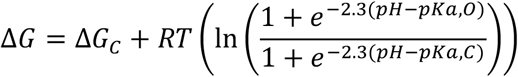

Because structural perturbations caused by ionization of internal Lys residues do not affect Trp. fluorescence, the closed and open states observed in this study cannot be distinguished in a chemical denaturation experiment, therefore the ΔG represents the simulated thermodynamic stability of the protein as observed in a chemical denaturation experiment. For the purpose of the simulation (solid gray curve in Figure 2D), the initial stability (set as the maximum stability of the closed state, ΔG_C_) was set to 8 kcal/mol. The terms p*K*_a,O_ and p*K*_a,C_ refer to the fully unfolded state p*K*_a_ value of the Lys residue (set to 10.2) and the apparent p*K*_a_ value of its titration when the protein is folded (set to 7.1), respectively. For the simulation of the closed state (dashed blue line in Figure 2D), the maximum stability was set to 8 kcal/mol, and the terms p*K*_a,C_ and p*K*_a,O_ refer to the pKa values in closed (set to 4 in this simulation, much lower than p*K*_a_^App^) and open (set to 10.2) states. Since the p*K*_a_ value of the Lys residue in the open state is similar to that of the Lys in water, its stability is not pH dependent and was set to reflect the loss in the stability of the closed state necessary to shift the p*K*_a_ value of the Lys from the unfolded state to its 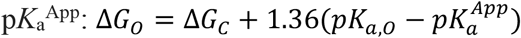

### Measurement and analysis of thermodynamic parameters

The values for ΔG_O_ were obtained from a previous study on the pH-dependence of the folding stability of these variants^2^, measurements and errors of which are available in Table S4. In general, the errors for stability measurements was between 0.1-0.2 kcal/mol, For the ΔG_O_ term, ΔG°_H2O_ values were chosen that were 0.5-1 pH unit below the apparent p*K*_a_ value of the Lys residue, unless the variant showed a second region of pH dependence in its thermodynamic stability after the Lys is protonated (indicative of the a p*K*_a_ shift of a surface residue^2^). In these cases, a ΔG°_H2O_ value 0-0.5 pH units below the apparent p*K*_a_ value were chosen. To estimate the population of partially folded state in the native state ensemble, the energy difference between the open state and the background at equivalent solution conditions was used to calculate a relative population of open versus native protein, using the following relation:

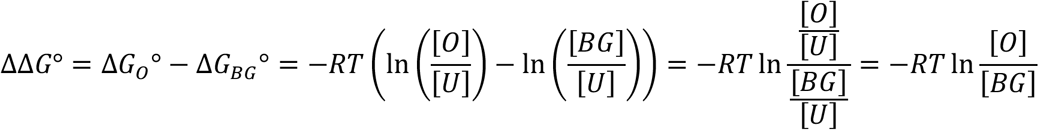

It is important to note that this analysis is solely meant to provide an estimate of populations, as the substitution itself could lead to favorable or unfavorable interactions that influence the energy differences between states. These are not likely to significantly shift the order of magnitude of the populations, as the energy landscape described by charge burial agrees well with that described by hydrogen deuterium exchange studies. It is also important to note that in many cases, different Lys residues that disrupt similar structural elements of the protein are not likely to share similar environments in their open states, as in many cases they are on opposing elements of secondary structure. This is true for variants in the black, green, and blue clusters in Figure 3, as well as to a lesser extent the yellow cluster (i.e., variants with K34 and K90, K66 and K25, K66 and K99, or K104 and K125); nevertheless, these cases have similar open state stabilities (within 1 kcal/mol of each other), again suggesting that the populations obtained from the thermodynamic analysis are indeed relevant to the native state ensemble of these proteins.

